# *Toxoplasma gondii* AP2IX-4 regulates gene expression during bradyzoite development

**DOI:** 10.1101/104208

**Authors:** Sherri Huang, Michael J. Holmes, Joshua B. Radke, Dong-Pyo Hong, Ting-Kai Liu, Michael W. White, William J. Sullivan

## Abstract

*Toxoplasma gondii* is a protozoan parasite of great importance to human and animal health. In the host, this obligate intracellular parasite persists as a tissue cyst form that is invisible to the immune response and unaffected by current therapies. The tissue cysts facilitate transmission through predation and give rise to chronic cycles of toxoplasmosis in immune compromised patients. Transcriptional changes accompany conversion of the rapidly replicating tachyzoites into the encysted bradyzoites, yet the mechanisms underlying these alterations in gene expression are not well-defined. Here we show that AP2IX-4 is a nuclear protein exclusively expressed in tachyzoites and bradyzoites undergoing division. Knockout of AP2IX-4 had no discernible effect on tachyzoite replication, but resulted in a reduced frequency of tissue cyst formation following alkaline stress induction – a defect that is reversible by complementation. AP2IX-4 has a complex role in regulating bradyzoite gene expression, as many bradyzoite mRNAs dramatically increased beyond normal stress-induction in AP2IX-4 knockout parasites exposed to alkaline media. The loss of AP2IX-4 also resulted in a modest virulence defect and reduced cyst burden in chronically infected mice, which was also reversed by complementation. These findings illustrate that the transcriptional mechanisms responsible for tissue cyst development operate across the intermediate life cycle from the dividing tachyzoite to the dormant bradyzoite.

## IMPORTANCE

*Toxoplasma gondii* is a single-celled parasite that persists in its host as a transmissible tissue cyst. How the parasite converts from its replicative form to the bradyzoites housed in tissue cysts is not well understood, but clearly involves changes in gene expression. Here we report that parasites lacking a cell cycle-regulated transcription factor called AP2IX-4 display reduced frequencies of tissue cyst formation in culture and in a mouse model of infection. Parasites missing AP2IX-4 lose the ability to regulate bradyzoite genes during tissue cyst development. Expressed in developing bradyzoites still undergoing division, AP2IX-4 may serve as a useful marker in the study of transitional forms of the parasite.

## INTRODUCTION

*Toxoplasma gondii* is an obligate intracellular protozoan parasite that has infected up to one third of the world’s population (1). A key factor contributing to this parasite’s widespread seroprevalence is its ability to transmit to new hosts through multiple routes (2). Felines are the definitive hosts that release infectious oocysts into the environment following infection. Oocysts, which are stable in the environment for up to one year, can be picked up by any warm-blooded vertebrate from contaminated soil or water. Upon ingestion, the sporozoites inside the oocysts are released, developing into tachyzoites capable of infecting any nucleated cell. During an initial infection, *Toxoplasma* is also capable of crossing the placenta and potentially infecting the unborn fetus, resulting in spontaneous abortion or birth defects (3). Tachyzoites replicate asexually and can lead to the destruction of the host cell; alternatively, tachyzoites can differentiate into bradyzoites. Bradyzoite development is accompanied by slowed parasite growth and the formation of tissue cysts that can persist for long periods, perhaps for the lifetime of the host. New hosts can become infected when these tissue cysts are consumed through predation, which includes humans who eat raw or undercooked meat containing tissue cysts. Life-threatening episodes of acute toxoplasmosis most commonly arise in patients with compromised immune systems, such as in HIV/AIDS, cancer chemotherapy, or organ transplantation as a result of newly acquired or recrudescent infection (4). A better understanding of the molecular mechanisms that drive interconversion between tachyzoites and bradyzoites is required to manage transmission and pathogenesis of *Toxoplasma*.

Many studies have documented genes that are exclusively expressed in either tachyzoites or bradyzoites, suggesting that gene expression mechanisms play a major role in coordinating developmental transitions in *Toxoplasma*. Studies of stage-specific promoters revealed that conventional *cis*-acting mechanisms operate to regulate developmental gene expression during tissue cyst formation (5). The principle class of transcription factor likely to work through these *cis*-acting promoter elements shares features with the Apetala-2 (AP2) family in plants (6). In plants, the AP2 domain is composed of a ~60 amino acid DNA-binding domain; plant factors with a single AP2 domain are linked to stress responses whereas those with tandem AP2 domains are linked to plant development (7). However, more recent analyses of AP2 domains have been performed on additional species, including prokaryotes and free-living photosynthetic algae closely related to apicomplexan parasites. These data suggest that apicomplexan AP2 (ApiAP2) proteins originated in bacteria and expanded in myzozoans independent of those in plants, prior to the split between the Apicomplexa and chromerids (8).

Numerous ApiAP2 factors found in *Plasmodium* spp., *Toxoplasma*, *Cryptosporidium parvum*, and *Theileria annulata* are capable of binding specific DNA motifs (9–12), and several have been implicated as transcription factors involved in stage-specific gene regulation. In the rodent malaria parasite, *Plasmodium berghei*, AP2-O activates gene expression in ookinetes (13), AP2-Sp regulates gene expression during the sporozoite stage (14), and AP2-L contributes to liver-stage development (15). PbAP2-G and PbAP2-G2 appear to work in conjunction to orchestrate transition of asexual blood-stage parasites to gametocytes (16). While PbAP2-G2 represses genes that are required for the proliferation of the asexual stage, PbAP2-G, and its orthologue in *P. falciparum*, function as a master regulator that is essential for sexual stage development (17, 18). These studies support that transcriptional switches underlie developmental transitions between parasite life cycle stages. This work has also supported a stochastic model for gametocyte development, whereby spontaneous activation of PfAP2-G in some asexual stage parasites prompts transition to the sexual stage (18).

There are 67 predicted proteins in the *Toxoplasma* genome that harbor one or more AP2 domains (see ToxoDB.org), more than double the number found in *Plasmodium* spp. (19). In *Toxoplasma*, ApiAP2 factors have been found in association with chromatin remodeling machinery (20), and genetic studies have supported roles in regulating transcription in the context of developmental transitions from tachyzoites to bradyzoites (21-23). AP2IX-9, which is a stress-induced ApiAP2 factor, binds to *cis*-regulatory elements of bradyzoite promoters and operates as a repressor of bradyzoite development (21). In contrast, the stress-induced factor, AP2IV-3, is an activator of bradyzoite gene expression and parasites lacking this factor display a reduced capacity to form tissue cysts *in vitro* (24). A knockout of AP2XI-4 impaired the activation of genes expressed during stress-induced differentiation and resulted in reduced cyst burdens in infected mice (22). Finally, AP2XII-6 was disrupted in a bradyzoite differentiation mutant generated by insertional mutagenesis (23). To date, no master ApiAP2 factor has been identified in *Toxoplasma* that drives the unique transcriptome of any life cycle stage.

Here, we investigated the function of AP2IX-4, an ApiAP2 whose mRNA is upregulated during bradyzoite differentiation and exhibits cell cycle regulation. We show that AP2IX-4 protein is expressed in the parasite nucleus during the S/M phase. Surprisingly, genetic ablation of AP2IX-4 has no discernible effects on tachyzoites, yet the knockout shows a reduced capacity to form tissue cysts *in vitro* and *in vivo*. These defects in developmental competency are reversed in a complemented knockout. AP2IX-4 is expressed in a subpopulation of bradyzoites that are undergoing division within a tissue cyst and acts to repress bradyzoite-associated genes. These findings underscore the complexity of the transcriptional program driving developmental transition to the bradyzoite stage and suggest that AP2IX-4 helps control the competing needs of parasite replication with formation of tissue cysts.

## RESULTS

### AP2IX-4 is expressed in the nucleus during the S-M phase of the cell cycle

A previous study of predicted *Toxoplasma* ApiAP2 proteins revealed that AP2IX-4 (TGME49_288950) is 1 of 7 ApiAP2 genes whose mRNA is up-regulated during alkaline stress (21) and exhibits cell cycle regulation in the tachyzoite (19). AP2IX-4 mRNA is one of the few ApiAP2 factors expressed at higher levels in day 21 tissue cysts purified from mice, and it is also robustly expressed during sporozoite meiosis (24). The single exon AP2IX-4 gene is predicted to encode a 104 kDa protein consisting of 951 amino acids that harbors a single AP2 domain; no other protein domains were detected with SMART or Interpro protein sequence analyzers (25, 26).

We generated an endogenously epitope-tagged version of AP2IX-4 to confirm its size and expression pattern at the protein level, and also to determine protein localization. RH*Δku80*::HXGPRT (RHQ) parasites were engineered to express native AP2IX-4 fused to a hemagglutinin (3xHA) tag at its C-terminus (AP2IX-4^HA^) (Fig. 1A). Western blotting of tachyzoite lysate shows that AP2IX-4^HA^ migrates as a single band at the expected size (Fig. 1B). Immunofluorescence assays (IFAs) were performed to detect subcellular localization throughout the tachyzoite cell cycle, co-staining with Inner Membrane Complex-3 (IMC3) antibody to help delineate cell cycle stages (27). AP2IX-4^HA^ protein was undetectable in the G1 phase, appeared during the S-M stages, then decreased in expression during cytokinesis (Fig. 1C), consistent with its reported mRNA profile (19). Since the IMC3 staining pattern does not distinguish between the G1 and S phases of the cell cycle, we quantified the proportion of vacuoles in G1/S with no HA signal to those in G1/S with HA signal and M/C with HA signal. Findings show the proportions to be approximately 60%, 30%, and 10%, which is consistent with the distribution of G1, S, and M/C cell cycle phases in an asynchronous tachyzoite population (Fig. 1D) (28). The IFAs also indicate that AP2IX-4^HA^ is exclusively present in the parasite nucleus when expressed (Fig. 1C).

**Figure 1.**
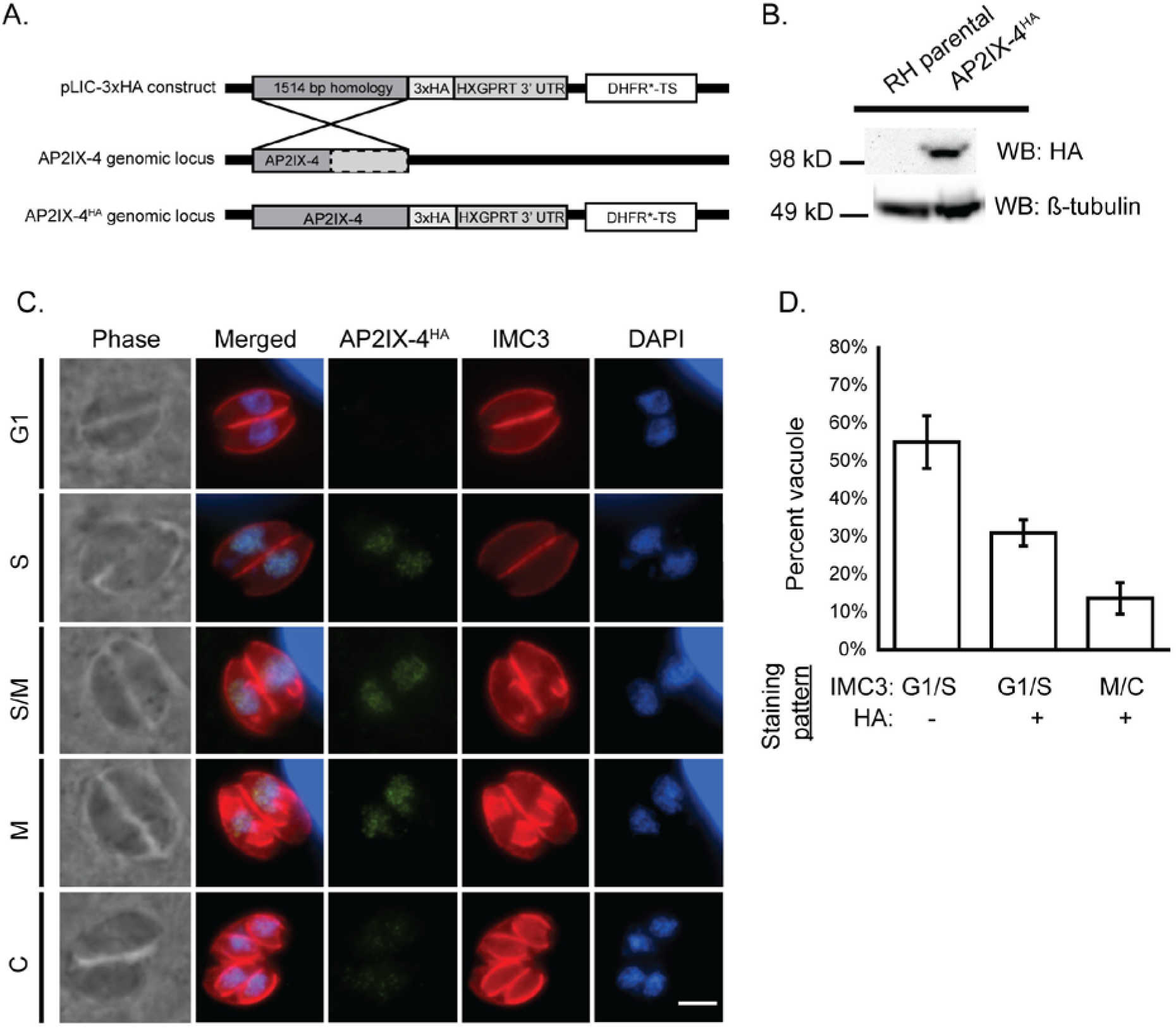
Characterization of AP2IX-4 protein expression. A. Schematic showing the strategy used to generate parasites expressing AP2IX-4 tagged at the C-terminus with 3xHA epitopes (AP2IX-4^HA^) in RHQ parasites. The AP2IX-4 genomic locus is aligned with the pLIC-3xHA construct used for endogenous tagging via single homologous recombination. The construct contains a 1,514 bp homology region and a DHFR*-TS drug selection cassette. B. Western blot of parental RHQ and AP2IX-4^HA^ parasites probed with anti-HA antibody. Anti-β-tubulin was used to verify presence of parasite protein. kD = kilodaltons. C. IFAs performed on AP2IX-4^HA^ tachyzoites using anti-HA (green). To monitor cell cycle phases, parasites were co-stained with anti-IMC3 (red) and DAPI (blue). G1, gap phase; S, synthesis phase; M, mitotic phase; C, cytokinesis. Scale bar = 3 microns. D. Quantification of the proportion of parasite vacuoles (out of 100) in the asynchronous population of AP2IX-4^HA^ parasites exhibiting G1/S IMC3 staining with absence of HA signal (G1/S HA-), G1/S IMC3 staining with positive HA signal (G1/S HA+), or M/C staining with positive HA signal. Error bars represent standard deviation from 3 independent experiments.

### AP2IX-4 is dispensable for tachyzoite replication *in vitro*

To investigate the function of AP2IX-4, we generated gene knockouts in RHQ and Pru*Δku80Δhxgprt* (PruQ) strains by replacing the genomic CDS with a selectable marker (DHFR*-TS) via double homologous recombination (Fig. 2A). Genomic PCRs were performed to screen transfected clones isolated by limiting dilution. Figure 2B shows the results confirming successful allelic replacement of the *AP2IX-4* gene in the PruQ background (named *Δap2IX-4*). RT-PCR was performed on mRNA harvested from *Δap2IX-4* parasites to verify loss of transcript (Fig. 2B). Parasite replication rates were determined using a doubling assay (Fig. 2C) and viability was examined using a plaque assay (Fig. 2D). Neither assay revealed a significant difference in the *in vitro* growth between parental and *Δap2IX-4* parasites in PruQ strain (Fig. 2C-D) or RHQ strain (data not shown). These results suggest that, despite its appearance during the S-M stages of the cell cycle, AP2IX-4 is not required for tachyzoite replication.

**Figure 2.**
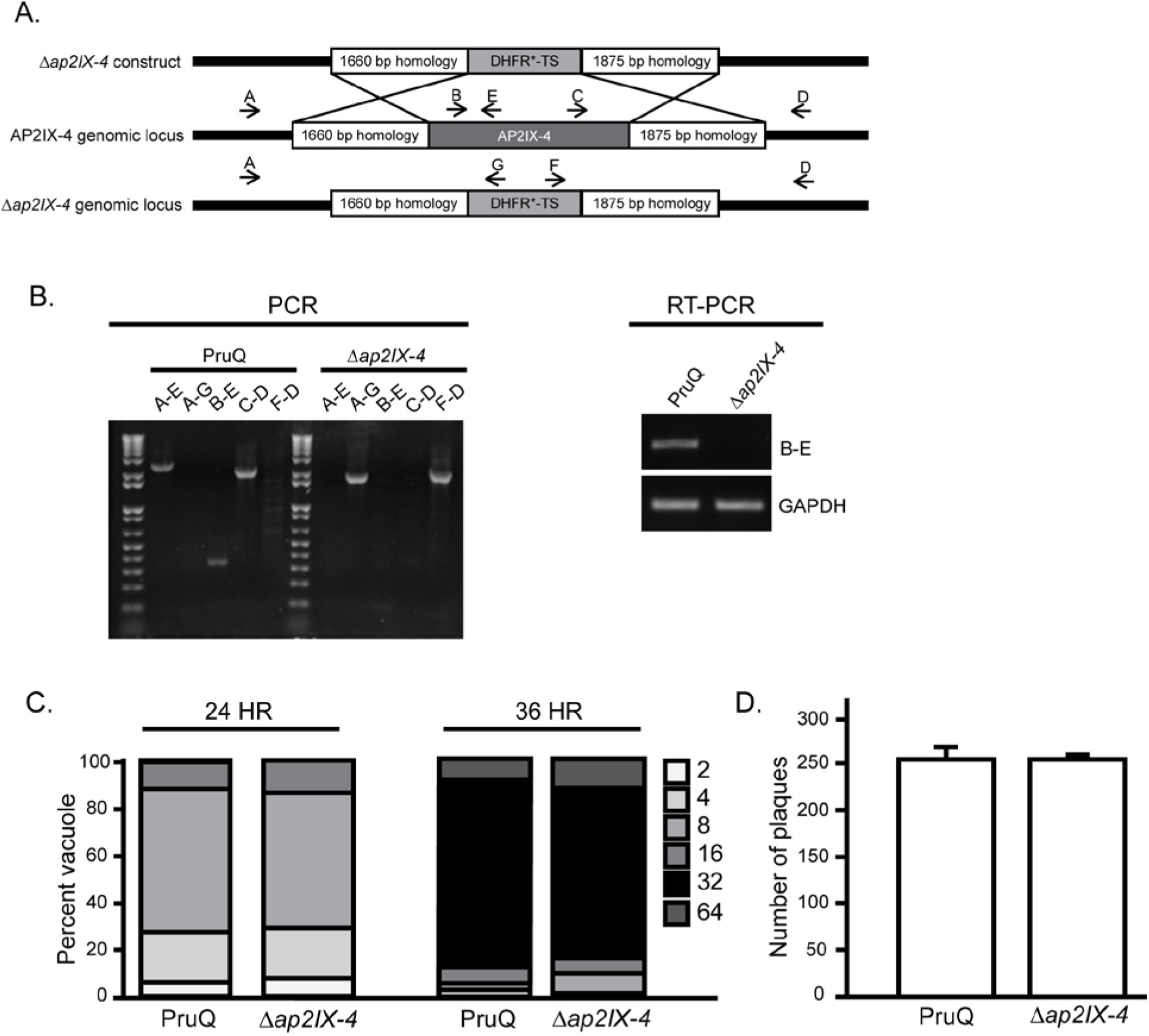
Generation of PruQ*Δap2IX-4* parasites. A. Schematic illustrating the generation of the AP2IX-4 knockout by allelic replacement with a DHFR*-TS minigene. B. Genomic PCRs were performed with the indicated primers (A-G) to verify replacement of the AP2IX-4 genomic locus with the DHFR*-TS minigene. RT-PCR with primers B and E confirmed the absence of AP2IX-4 transcripts in PruQ*Δap2IX-4* parasites. Primers for GAPDH were used as a positive control for the RNA preparation. C. Representative doubling assay showing the number of parasites/vacuole at 24- and 36-hour time-points post-infection for parental PruQ versus PruQ*Δap2IX-4* parasites. Three independent assays were performed with similar results. D. Plaque assays were performed to compare *in vitro* viability of PruQ versus PruQ*Δap2IX-4* parasites. Three independent assays were performed with similar results.

### AP2IX-4 is present in bradyzoites and required for efficient cyst formation *in vitro*

Previous studies indicate that AP2IX-4 mRNA increases during bradyzoite differentiation *in vitro* and that elevated levels of AP2IX-4 transcripts are present during the chronic stage of infection in mice (19, 29). To examine the function of AP2IX-4 in bradyzoites, we complemented the PruQ*Δap2IX-4* parasites by targeting a cDNA-derived version of AP2IX-4 tagged with HA at its C-terminus to be under control of the native AP2IX-4 promoter (PruQ*Δap2IX-4*::AP2IX-4^HA^, Fig. 3A). Genomic PCRs confirmed integration of the complementation construct at the disrupted *ap2IX-4* locus (Fig. 3B) and Western blot analysis confirms the expected size of the AP2IX-4^HA^ protein (Fig. 3C). Furthermore, the AP2IX-4^HA^ protein in the complemented knockout localizes properly to the parasite nucleus during S-M phases (Fig. 3D), matching the pattern we observed for native protein in RH parasites (Fig. 1C). The proportion of vacuoles in G1/S with no HA signal to those in G1/S with HA signal and M/C with HA signal were approximately 60%, 30%, and 10% (Fig. 3E), consistent with observations made in type I strain (Fig. 1D). Together, these data confirm the fidelity of the complemented knockout and uncover no differences in localization or cell cycle regulation of AP2IX-4 between type I and type II strains.

**Figure 3.**
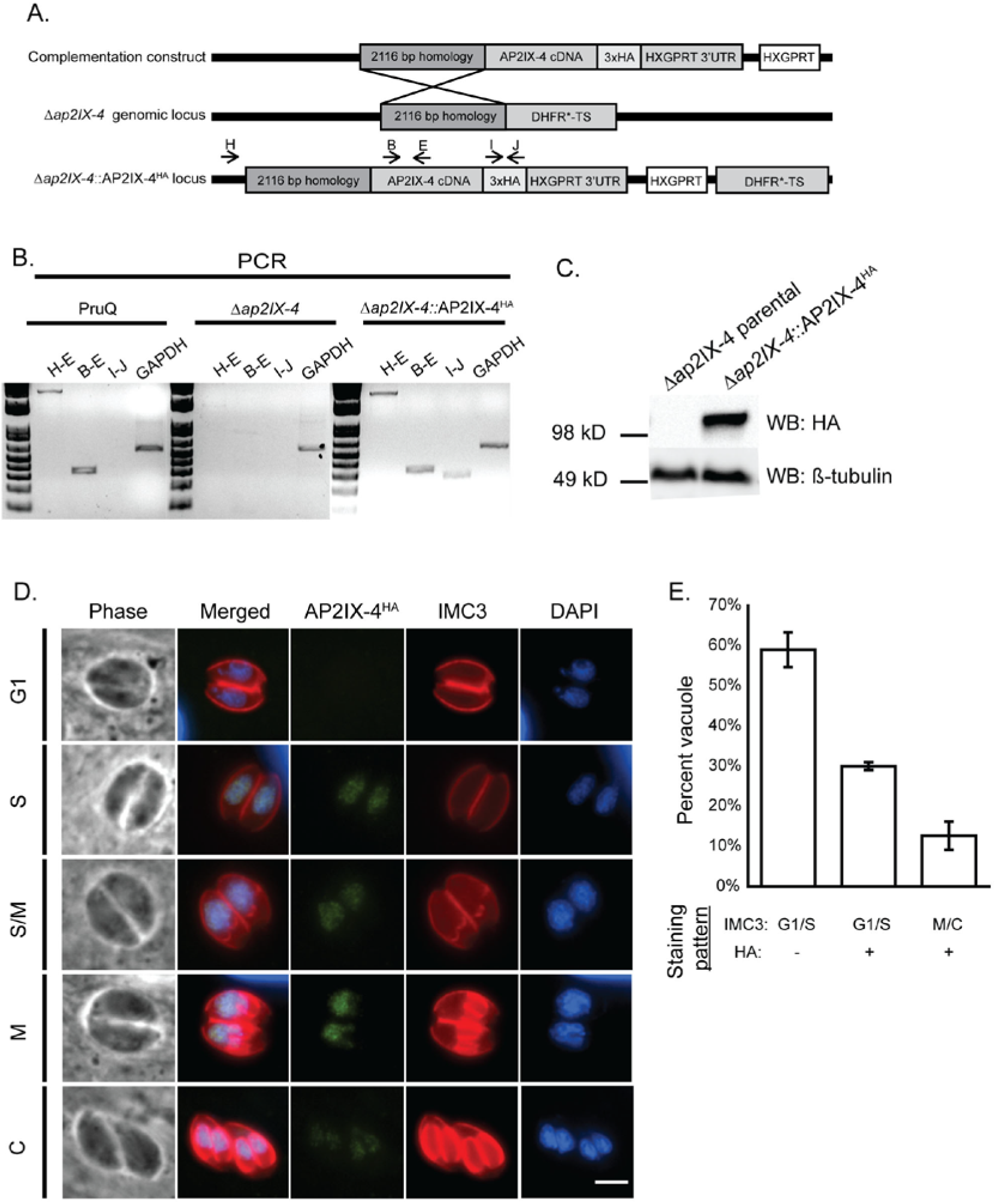
Complementation of the AP2IX-4 knockout. A. Schematic illustrating the generation of the complemented knockout (*Δap2IX-4*::AP2IX-4^HA^). A cDNA-derived version of AP2IX-4 tagged at the C-terminus with 3xHA (complementation construct) was targeted to the ablated genomic locus in *Δap2IX-4* parasites. B. Genomic PCRs with indicated primer pairs were performed to confirm proper integration of the complementation construct into *Δap2IX-4* parasites. Primers to GAPDH were used as a positive control of the DNA preparation. C. Western blot of *Δap2IX-4* and *Δap2IX-4*::AP2IX-4^HA^ parasites with anti-HA, using anti-β-tubulin as a control to verify the presence of parasite protein. D. IFAs were performed on *Δap2IX-4*::AP2IX-4^HA^ tachyzoites as described in Figure 1C. E. Quantification of the proportion of parasite vacuoles (out of 100) in the asynchronous population of AP2IX-4^HA^ parasites performed as described in Figure 1D.

Taking advantage of this HA-tagged version of AP2IX-4 in the developmentally competent type II strain, we monitored protein expression by IFA during alkaline stress (pH 8.2), a well-characterized inducer of tissue cyst development *in vitro* (30). The PruQ strain also contains a green fluorescent protein (GFP) reporter driven by the bradyzoite-specific LDH2 promoter (31, 32). Under alkaline stress bradyzoite induction conditions, expression of AP2IX-4^HA^ is heterogeneous within the population of bradyzoites housed in individual cysts (Fig. 4A). Comparing the number of AP2IX-4^HA^–positive parasites to the total number of parasites within developing tissue cysts (among 50 random cysts in 3 independent replicates), we found that only 13% are expressing AP2IX-4^HA^ at 2 and 4 days post-induction. In contrast, 37% of tachyzoites express AP2IX-4^HA^. Co-staining for IMC3 reveals that GFP-positive bradyzoites expressing AP2IX-4^HA^ are in the process of dividing, revealing that AP2IX-4^HA^ expression continues to be cell cycle-regulated during encystation (Fig. 4B). Loss of AP2IX-4 does not appear to have a significant impact on the number of parasites undergoing division at day 4 of bradyzoite induction; the proportion of dividing parasites as assessed by IMC3 staining of 50 random cysts was 7.6% for PruQ parasites and 6.5% for PruQ*Δap2IX-4* parasites.

**Figure 4.**
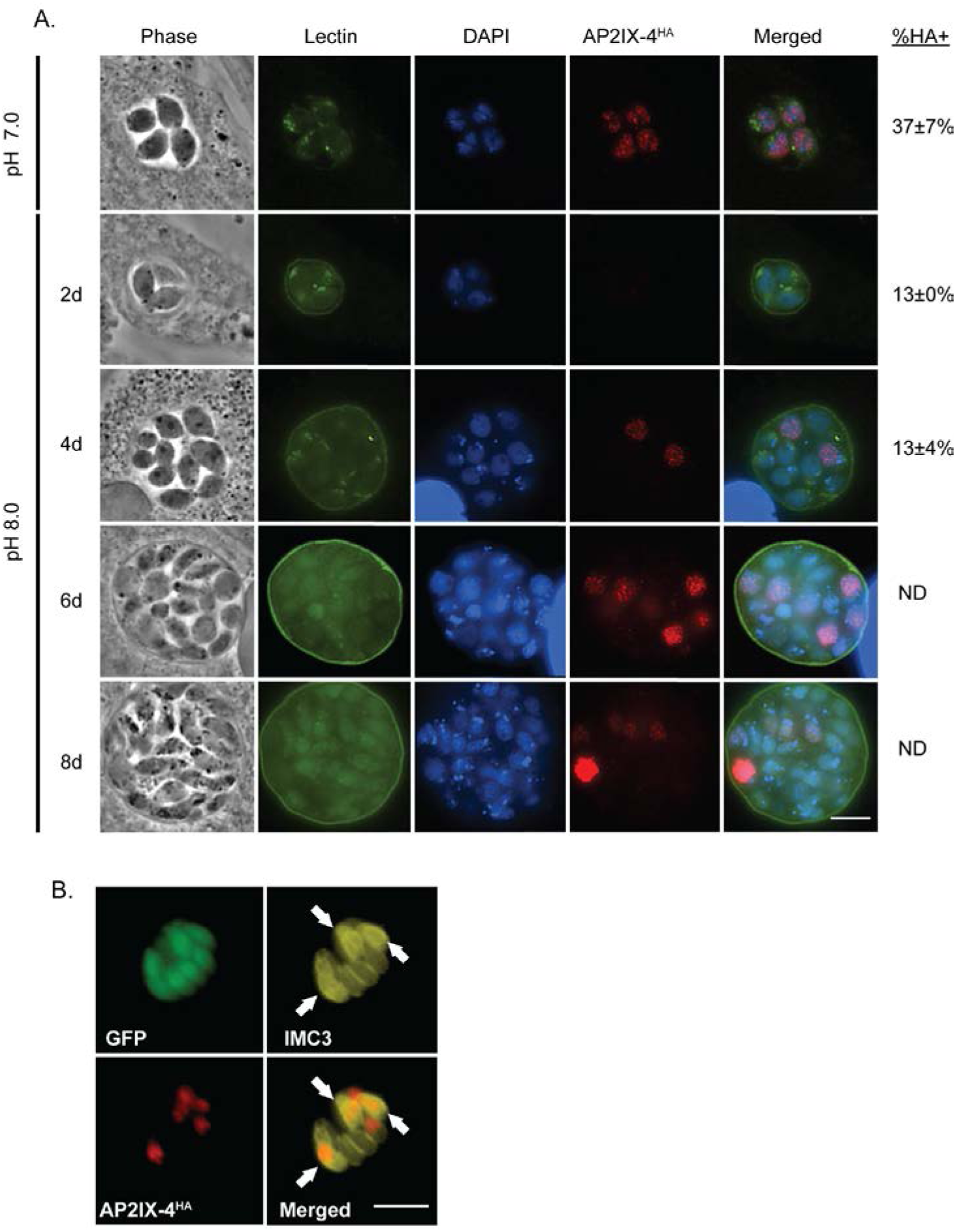
Expression of AP2IX-4 in developing bradyzoites *in vitro*. A. IFAs were performed on *Δap2IX-4*::AP2IX-4^HA^ parasites during tachyzoite and developing bradyzoite stages up to 8 days in alkaline pH. FITC-conjugated *Dolichos* lectin was used to visualize bradyzoite cyst walls (the LDH2-GFP reporter also appears in bradyzoites in this channel). AP2IX-4^HA^ protein was visualized using anti-HA (red) and DAPI (blue) was used to stain DNA. The percent of parasites expressing AP2IX-4^HA^ within 50 random vacuoles was recorded (%HA+), noting that it was not possible to accurately discern the total number of parasites in tissue cysts beyond 4 days post-induction (ND = not determined). Values represent the average and standard deviation for 3 independent experiments. B. IFAs performed with anti-IMC3 antibody (yellow) on *Δap2IX-4*::AP2IX-4^HA^parasites following 4 days in alkaline pH. The LDH2-GFP reporter (green) identifies bradyzoites. AP2IX-4^HA^ protein was visualized using anti-HA (red). Arrowheads point to budding daughter cells in dividing parasites expressing AP2IX-4^HA^. Scale bar = 3 microns.

We next examined the frequency of bradyzoite cyst development in parental PruQ and PruQ*Δap2IX-4* parasites in response to alkaline stress, as determined by the number of vacuoles staining positive for cyst wall protein binding *Dolichos* lectin. After 4 days in alkaline pH, 73% of parental PruQ parasites converted into bradyzoites, but this was reduced to 46% in parasites lacking AP2IX-4 (Fig. 5). Complementation of the AP2IX-4 knockout parasites restored the rate of bradyzoite cyst formation to levels matching wild-type PruQ strain (Fig. 5). These data show that AP2IX-4^HA^ continues to be expressed in a cell cycle-regulated manner post-bradyzoite induction and its ablation impairs the frequency at which cysts form *in vitro*.

**Figure 5.**
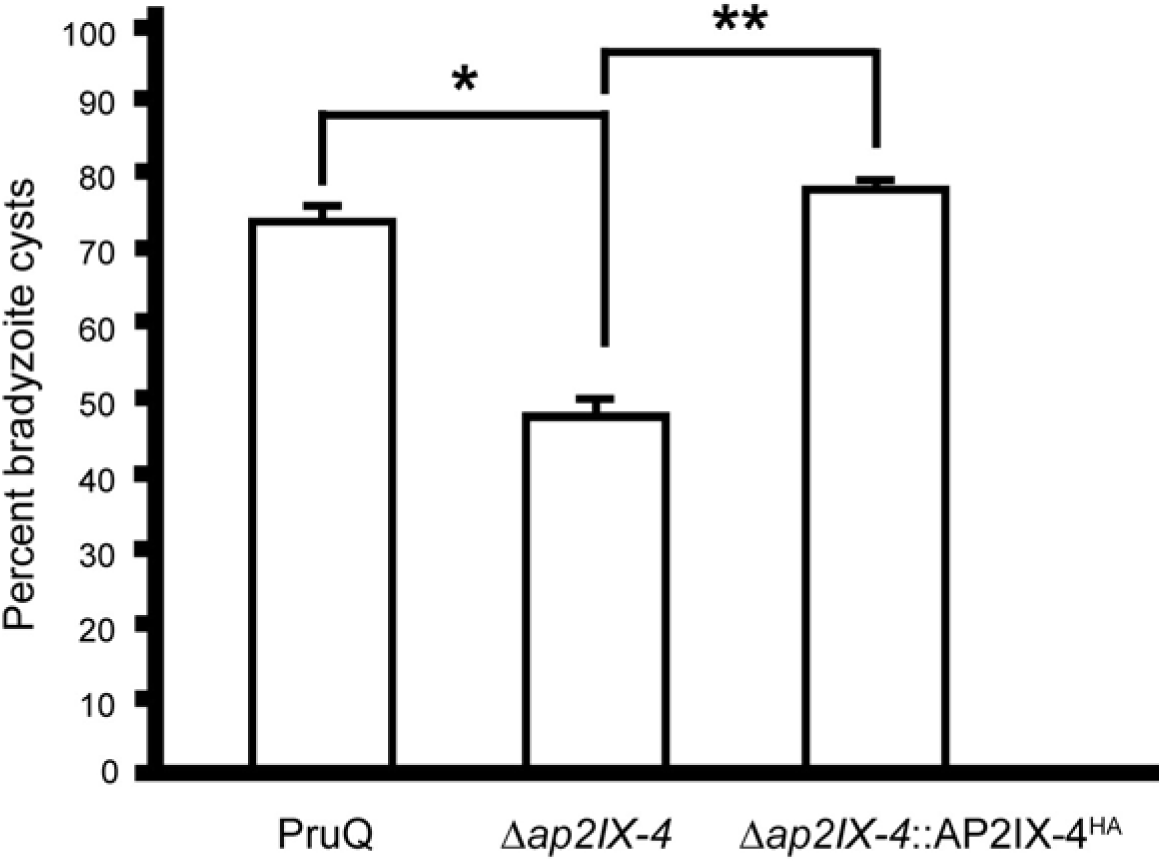
Loss of AP2IX-4 results in lower frequency of bradyzoite cyst formation *in vitro*. Parental PruQ, *Δap2IX-4*, and *Δap2IX-4*::AP2IX-4^HA^ parasites were cultured in alkaline medium for 4 days and then stained with *Dolichos* lectin to visualize tissue cyst walls. For each sample, 100 random vacuoles were surveyed for the presence or absence of lectin staining to determine frequency of cyst formation. N=4 for parental and *Δap2IX-4*; N=3 for *Δap2IX-4*::AP2IX-4^HA^. * denotes p=0.04, ** denotes p=0.006, unpaired two-tailed Student’s t-test. Error bars represent standard error of the mean.

### Loss of AP2IX-4 results in dysregulated gene expression

To assess the impact of AP2IX-4 on gene expression, RNA was harvested from parental PruQ and PruQ*Δap2IX-4* parasites cultured under tachyzoite (pH 7) or bradyzoite-inducing (pH 8.2) conditions for 2 days and hybridized onto ToxoGeneChip microarrays (33). Under tachyzoite conditions, very few genes were dysregulated in the knockout (Supplemental Table 2). In contrast to the tachyzoite culture conditions, 119 genes were differentially expressed >2-fold in the knockout during bradyzoite induction conditions: 64 were downregulated in the PruQ*Δap2IX-4* compared to the parental while 55 were upregulated (Supplemental Table 3). Approximately 89% of these genes were complemented in knockout parasites in which AP2IX-4 expression had been restored. Table 1 shows a representative group of genes that were rescued by complementation (the complete list of complemented genes is in Supplemental Table 3).

Four bradyzoite-associated genes were downregulated in the knockout following stress including enoyl coA hydratase/isomerase, and hypothetical proteins TGME49_205680, TGME49_287040, and TGME49_306270 (Table 1). Enoyl coA hydratase/isomerase, along with a MoeA domain-containing protein (involved in biosynthesis of molybdopterin), were downregulated in the AP2IX-4 knockout in both tachyzoite and bradyzoite-inducing conditions. Strikingly, the majority of genes showing enhanced expression in the AP2IX-4 knockout upon stress were those encoding well-established bradyzoite-associated proteins such as ENO1, BAG1, PMA1, B-NTPase, LDH2, DnaK-TPR, SAG2C, and MCP4 (Table 1).

**Table 1.** Differentially expressed genes (FC≥2) in *∆ap2IX-4* and *∆ap2IX-4*::AP2IX-4^HA^ during bradyzoite induction conditions. Genes that were previously found to be upregulated during bradyzoite induction are highlighted in gray (see Materials and Methods).

| Accession no. | Annotation | PruQ$\Delta ap2IX-4$<br>vs. PruQ parental<br>(FC) | $\Delta ap2IX-4::AP2IX-4^{HA}$<br>vs PruQ$\Delta ap2IX-4$<br>(FC) |
| --- | --- | --- | --- |
| **DOWNREGULATED GENES** |  |  |  |
| TGME49_288950 | AP2IX-4 | -38.1 | 22.00 |
| TGME49_319890 | hypothetical protein | -7.6 | 1.8 |
| TGME49_293480 | MoeA N-terminal region (domain I and II) domain-containing protein | -7.4 | 3.4 |
| TGME49_243940 | hypothetical protein | -7.0 | 1.8 |
| TGME49_204050 | subtilisin SUB1 (SUB1) | -4.9 | 1.7 |
| TGME49_317705 | enoyl coA hydratase/isomerase | -3.8 | 2.2 |
| TGME49_297910 | hypothetical protein | -3.7 | 2.1 |
| TGME49_256792 | hypothetical protein | -3.6 | 2.1 |
| TGME49_217400 | hypothetical protein | -3.5 | 3.2 |
| TGME49_205680 | hypothetical protein | -3.2 | 1.7 |
| TGME49_287040 | hypothetical protein | -3.1 | 1.8 |
| TGME49_306270 | hypothetical protein | -3.0 | 1.7 |
| TGME49_280500 | inorganic anion transporter, sulfate permease (SulP)<br>family protein | -2.6 | 2.2 |
| TGME49_261250 | histone H2A1 | -2.6 | 1.7 |
| TGME49_240090 | roptry kinase family protein ROP34, putative | -2.5 | 1.6 |
| TGME49_214190 | SRS46 | -2.3 | 1.6 |
| TGME49_239260 | histone H4 | -2.2 | 1.7 |
| TGME49_261240 | histone H3 | -2.1 | 2.4 |
| TGME49_305160 | histone H2Ba | -2.0 | 2.9 |
| TGME49_255060 | cytochrome b(N-terminal)/b6/petB subfamily protein | -2.0 | 3.9 |
| **UPREGULATED GENES** |  |  |  |
| TGME49_268860 | ENO1 | 10.2 | -2.3 |
| TGME49_312905 | hypothetical protein | 9.7 | -7.3 |
| TGME49_209755 | hypothetical protein | 7.0 | -2.5 |
| TGME49_259020 | BAG1 | 5.7 | -1.6 |
| TGME49_293790 | hypothetical protein | 4.9 | -2.1 |
| TGME49_278080 | Toxoplasma gondii family A protein | 4.9 | -1.7 |
| TGME49_216140 | tetratricopeptide repeat-containing protein | 4.4 | -1.8 |
| TGME49_252640 | P-type ATPase PMA1 | 4.4 | -1.7 |
| TGME49_225290 | GDA1/CD39 (nucleoside phosphatase) (B-NTPase) | 4.1 | -2.1 |
| TGME49_315560 | ATP-binding cassette G family transporter ABCG77 | 4.0 | -1.5 |
| TGME49_291040 | lactate dehydrogenase LDH2 (LDH2) | 4.0 | -1.6 |
| TGME49_202020 | DnaK-TPR | 3.9 | -1.8 |
| TGME49_221550 | CMGC kinase, MAPK family, MEK kinase-related | 3.7 | 1.9 |
| TGME49_292260 | SAG-related sequence SRS36B (SAG5D) | 3.6 | -1.9 |
| TGME49_266700 | hypothetical protein | 3.5 | -1.9 |
| TGME49_240470 | hypothetical protein | 3.2 | -1.7 |
| TGME49_207160 | SAG-related sequence SRS49D (SAG2C) | 3.1 | -2.2 |
| TGME49_267470 | cold-shock DNA-binding domain-containing protein | 3.1 | -1.7 |
| TGME49_253680 | hypothetical protein | 3.0 | 1.5 |
| TGME49_208730 | microneme protein, putative (MCP4) | 3.0 | -1.6 |
| TGME49_207150 | SAG-related sequence SRS49C (SAG2D) | 2.9 | -1.9 |
| TGME49_287960 | hypothetical protein | 2.7 | -1.6 |
| TGME49_218470 | protein disulfide-isomerase | 2.6 | 1.9 |
| TGME49_253710 | hypothetical protein | 2.6 | -1.7 |
| TGME49_306338 | dynein gamma chain, flagellar outer arm, putative | 2.5 | -1.9 |
| TGME49_243720 | peroxisomal biogenesis factor PEX11 (PEX11) | 2.5 | -1.9 |
| TGME49_259960 | nucleoside-diphosphatase | 2.4 | -1.5 |
| TGME49_312905 | hypothetical protein | 2.4 | -2.0 |
| TGME49_314250 | bradyzoite rhoptry protein BRP1 (BRP1) | 2.4 | -1.8 |
| TGME49_253330 | rhoptry kinase family protein (BPK1) | 2.3 | -1.5 |
| TGME49_224830 | hypothetical protein | 2.2 | 1.5 |
| TGME49_221840 | hypothetical protein | 2.2 | -1.5 |
| TGME49_311370 | methylmalonate-semialdehyde dehydrogenase | 2.1 | -2.0 |
| TGME49_213460 | hypothetical protein | 2.0 | -1.7 |
| TGME49_266900 | Cyclin, N-terminal domain-containing protein | 2.0 | -1.6 |

Findings from the transcriptomic analysis suggest that AP2IX-4 leads to aberrant gene expression that is more pronounced following alkaline stress treatment. The overall pattern that emerged suggests that bradyzoite-associated genes generally see enhanced activation in the knockout following stress. To determine if the dysregulated gene expression caused by ablation of AP2IX-4 has consequences on pathogenesis, we examined acute and chronic infection in a mouse model.

### Analysis of PruQ*Δap2IX-4* in mouse models of acute and chronic toxoplasmosis

To determine if the loss of AP2IX-4 impacts virulence *in vivo*, BALB/c mice (8 per group) were infected with 10^7^, 10^6^, or 10^5^ parental PruQ, PruQ*Δap2IX-4*, or PruQ*Δap2IX-4*::AP2IX-4^HA^ parasites and monitored for 14 days. A dose of 10^7^ parasites was equally lethal for all strains injected into the mice, whereas all mice infected with 10^5^ parasites survived (Fig. 6A). Three of the 8 infected mice infected with either parental or PruQ*Δap2IX-4*::AP2IX-4^HA^ parasites survived a dosage of 10^6^; however, all 8 mice infected with 10^6^ of the PruQ*Δap2IX-4* parasites survived (Fig. 6A). Despite being dispensable for tachyzoite replication *in vitro*, these data suggest that loss of AP2IX-4 results in a modest virulence defect *in vivo*.

**Figure 6.**
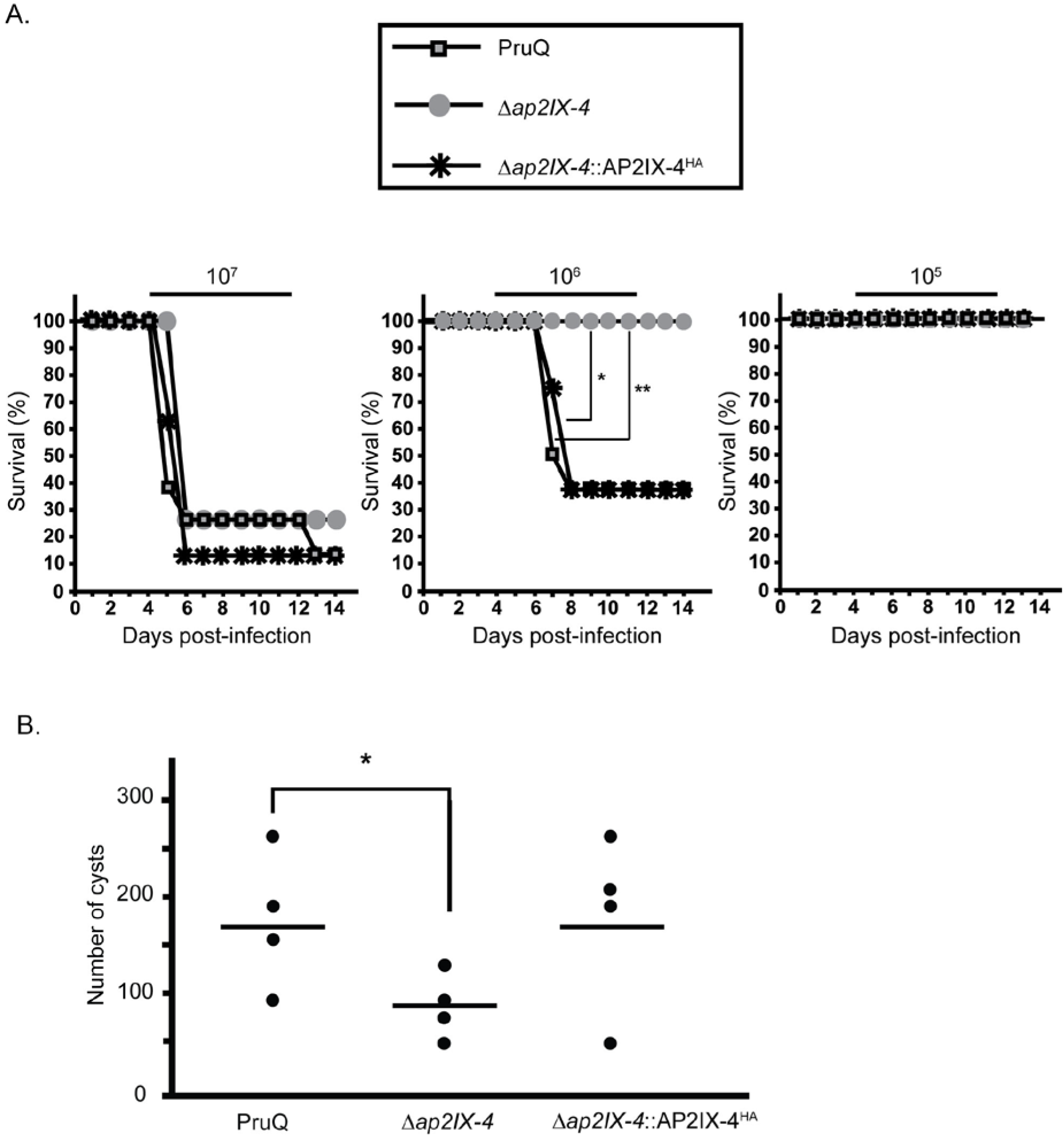
Virulence and cyst burden in mice infected with *Δap2IX-4* parasites. A. Mice (N=8 per group) were infected with either 10^7^, 10^6^, or 10^5^ parental, *Δap2IX-4*, or *Δap2IX-4*::AP2IX-4^HA^ parasites and monitored over 14 days for survival. * denotes p=0.01; ** denotes p=0.02, log-rank test. B. Mice were infected with 10^6^ of the designated parasites (N=4 per group) and allowed to progress to chronic infection for 35 days before brain cyst burden was assessed. * denotes p=0.03, unpaired one-tailed Student’s t-test.

We also analyzed the brain cyst burden in BALB/c mice infected with 10^6^ parental PruQ, PruQ*Δap2IX-4*, or PruQ*Δap2IX-4*::AP2IX-4^HA^ parasites at 35 days post-infection. Mice infected with parental parasites contained an average of 169 brain cysts while those infected with PruQ*Δap2IX-4* yielded a significantly reduced average of 91 brain cysts (Fig. 6B). Complementation of PruQ*Δap2IX-4* reversed the phenotype, producing an average of 176 brain cysts (Fig. 6B). This experiment was repeated to ensure reproducibility and yielded similar results (data not shown). Consistent with the reduced frequency of bradyzoite development noted *in vitro*, these data suggest that loss of AP2IX-4 also results in decreased bradyzoite frequency *in vivo*.

## DISCUSSION

The establishment of tissue cysts within host cells is essential for *Toxoplasma* transmission and is the underlying cause of chronic infection in patients. Tissue cyst formation requires a slowing of rapidly proliferating tachyzoites, which is associated with a lengthening of the G1 phase (34). A complete inhibition of tachyzoite growth blocks bradyzoite development (35), while no fast growing bradyzoite has ever been identified. These observations suggest that formation of bradyzoites is a step-wise process involving growth mechanisms, although how many steps and what molecular mechanisms are involved is not understood. At a minimum, tissue cysts development involves a slow growing pre-bradyzoite expressing bradyzoite antigens and a mature stage that is non-replicative and may be required for transmission (11, 36). Defining the bradyzoite developmental pathway is hampered by its heterologous nature. Unlike the tachyzoite stage, which shows striking synchronous intravacuolar behavior, parasites undergoing bradyzoite development are largely asynchronous and tissue cysts contain a highly variable number of bradyzoites (36). These challenges have contributed to the lack of mechanistic knowledge of how tissue cysts are formed.

Like all ApiAP2 factors characterized, AP2IX-4 is present exclusively in the nucleus. AP2IX-4 also belongs to the set of cyclically regulated ApiAP2 factors of the tachyzoite (Fig. 1C) (19). AP2IX-4 mRNA and protein are dynamically expressed in the tachyzoite with peak expression during the S phase and mitosis and cytokinesis; as newly formed tachyzoites emerge (G1 phase), AP2IX-4 expression falls below detectable limits.

To gain insight into the biological role of AP2IX-4, we generated knockout parasites in both Type I RH and Type II Pru strains. Surprisingly, AP2IX-4 is not essential to tachyzoite replication, indicating its role is not in regulating the tachyzoite cell cycle per se, however, this factor is required for efficient bradyzoite development *in vitro* and *in vivo*. These data indicate AP2IX-4 could be required for regulating the transition from the tachyzoite to bradyzoite. Previous studies have demonstrated that spontaneous bradyzoite gene expression occurs during the S/M phases of the tachyzoite cell cycle (19) and the pre-bradyzoite (SAG1+/BAG1+) transitional stages also show higher numbers of parasites with the characteristic genome content of late S phase/mitotic parasites (34). In response to alkaline-stress, AP2IX-4 expression was detected in only in a subpopulation of parasites (Fig. 4) that, based on co-staining with the daughter budding marker IMC3, were undergoing division. These AP2IX-4 vacuoles were also positive for the DBA staining indicating differentiation to the tissue cyst was also underway. These findings support the long held view that bradyzoite development is asynchronous compared to tachyzoite replication (36, 37). The stochastic nature of bradyzoite development is not understood mechanistically, however, ApiAP2 factors like AP2IX-4 now provide new experimental avenues to help define the individual steps of tissue cyst differentiation at the level of the individual parasite.

Transcriptome analysis of PruQ*Δap2IX-4* parasites shed further light on AP2IX-4 function. Consistent with a presumed lack of a role in tachyzoite cell division, no significant differences were determined between the Pru parent and AP2IX-4 knockout tachyzoites (grown in normal pH 7.0 media). However, in response to alkaline stress, numerous bradyzoite-specific mRNAs saw greatly enhanced expression when AP2IX-4 was disrupted (Table 1), with the vast majority of these gene expression changes (~90%) reversed following complementation. As only ~13% of encysted parasites are expressing AP2IX-4 (Fig. 4), it is likely that the lack of synchronization in the microarray study underestimated the magnitude and number of genes regulated by this factor. Nevertheless, our transcriptomic analysis indicates that AP2IX-4 primarily functions to repress a subset of bradyzoite genes during the early stages of bradyzoite development. In addition to metabolic genes known to be specifically expressed in the bradyzoite, AP2IX-4 may also control a novel, bradyzoite-specific CMGC (cyclin-dependent kinase [CDK], mitogen-activated protein kinase [MAPK], glycogen synthase kinase [GSK3], CDC-like kinase [CLK]) kinase that may play a role in stress-induced signal transduction promoting latency. Altogether, these data reveal that what is distinctive about AP2IX-4 is its role in balancing the induction of gene expression in stress-induced bradyzoites, while having no apparent function in regulating gene expression specific to the tachyzoite.

This pattern of gene expression in AP2IX-4 knockout parasites is not the classic profile of a transcriptional repressor, where mis-expression of bradyzoite genes in the tachyzoite would be expected. In a separate study, we have identified a bradyzoite transcriptional repressor in the tachyzoite that fits this classic pattern (38).

Like AP2IX-4, AP2IV-4 also exhibits peak expression in the tachyzoite S/M phase; however, dysregulated expression of bradyzoite-associated mRNAs in an AP2IV-4 knockout occurs without stress. By contrast, AP2IX-4 may act as a passive repressor (i.e. suppresses transcription by competing with an activator for its target binding site) that requires an additional protein induced by alkaline stress (39). Both types of repressors (active and passive) are well-documented among the AP2 transcription factors of plants (39). Alternatively, AP2IX-4 could be a weak transcriptional activator that works in conjunction with a stronger activator to express bradyzoite mRNAs at the proper level and time. In the absence of AP2IX-4 to regulate a stronger transcription factor, the timing of expression for a subset of bradyzoite-associated genes could become unbalanced, leading to negative consequences on tissue cyst development that we have observed *in vitro* and *in vivo*. In addition, the unbalancing of gene expression during bradyzoite development caused by the loss of AP2IX-4 could cause problems with either tissue-specific formation of the tissue cyst or in achieving the proper escape from the host immune system. Preventing the pre-arming of the immune system against the parasite surviving to form the tissue cyst is a confirmed function for AP2IV-4 (38) and likely has driven the evolution of AP2IX-4 function. Validating this function will require a complete dissection of the AP2IX-4 transcriptional mechanism and its molecular partners during bradyzoite development *in vitro* and *in vivo*.

Our recent studies, and other investigations, have uncovered a number of ApiAP2s that modulate gene expression associated with the developmental transition between tachyzoites and bradyzoites. The AP2IX-4 studied here joins this list of factors. AP2IX-9 represses the expression of bradyzoite mRNAs; overexpression of AP2IX-9 promotes tachyzoite growth during stress and prevents formation of tissue cysts (21). AP2IV-4 is a cell cycle-regulated factor with peak expression in the tachyzoite S/M phase that also suppresses bradyzoite gene expression (38). In contrast, AP2IV-3 is an activator of bradyzoite gene expression; a knockout reduces tissue cyst formation and its overexpression increases tissue cyst formation (24). Changes in bradyzoite gene expression also occurred following the deletion of AP2XI-4 suggesting this factor could be a transcriptional activator of bradyzoite gene expression (22). Thus, the model that is emerging from recent ApiAP2 factor studies is a complex transcriptional network of activators and repressors operating at the interface of tachyzoite replication and switching to the bradyzoite stage controls the intermediate life cycle. Multiple levels of ApiAP2 factor regulation in bradyzoite development contrasts with the "master regulatory factor" model of gametogenesis of *Plasmodium falciparum* (18). *Toxoplasma* has more than twice as many ApiAP2 factors as *P. falciparum* and a much larger number of hosts that can serve the *Toxoplasma* intermediate life cycle. Consistent with this life cycle flexibility, a recent analysis of ApiAP2 factor expression revealed that a larger number of ApiAP2 factors are expressed in the intermediate life cycle (24). Therefore, it is reasonable to suggest that the large diversity of *Toxoplasma* ApiAP2 factors regulating bradyzoite development was necessary to expand the host range of the intermediate life cycle. Future studies focused on how *Toxoplasma* ApiAP2 factors function in different animal hosts will be important to understand how these transcriptional mechanisms evolved.

## MATERIALS AND METHODS

### Parasite culture and growth assays

Parasites strains used in these studies include RH*Δku80*::HXGPRT (RHQ) and Pru*Δku80Δhxgprt* (PruQ) (31, 40, 41). All parasites were cultured in confluent monolayers of human foreskin fibroblasts (HFFs) using Dulbecco’s Modified Eagle’s Medium (DMEM) supplemented with 1% or 5% heat-inactivated fetal bovine serum (FBS), respectively. Cultures were maintained in humidified incubators at 37°C with 5% CO_2_. Parasite doubling assays were performed as previously described (42). Briefly, 10^6^ freshly lysed tachyzoites were inoculated onto host cell monolayers; after 24 hours, the number of tachyzoites present within 50 random vacuoles was recorded at 12 hour intervals. For plaque assays, infected monolayers in 12-well plates were stained with crystal violet to visualize and count plaques (43).

To initiate conversion to bradyzoites *in vitro*, type II parasites were allowed to invade HFF monolayers for 4 hours under normal culture conditions, then the medium was replaced with alkaline RPMI (pH 8.2) and the cultures moved to a 37°C incubator with ambient CO_2_.

### Generation of parasites expressing endogenously tagged AP2IX-4

RHQ parasites were engineered to express endogenous AP2IX-4 tagged at its C-terminus with three tandem copies of the hemagglutinin (3xHA) epitope. Modification of the endogenous genomic locus was achieved through allelic replacement using parasites lacking KU80 so to facilitate homologous recombination (40, 41). This genetic “knock-in” approach employed the pLIC-3xHA plasmid, which contains a 1,514 bp sequence from the AP2IX-4 locus for homologous recombination (Fig. 1), amplified with primers shown in Supplemental Table 1. This endogenous tagging construct also contains which contains a modified dihydrofolate reductase-thymidylate synthase (DHFR*-TS) minigene that confers resistance to pyrimethamine for selection of transgenic parasites (44). Following drug selection, parasites were cloned by limiting dilution in 96-well plates.

### Generation of AP2IX-4 knockout parasites

Deletion of the AP2IX-4 gene was carried out in both RHQ and PruQ parasites. To generate the *Δap2IX-4* clones, a plasmid was constructed that flanked the DHFR*-TS minigene with 1.6-1.9 kb sequences homologous to regions of the AP2IX-4 locus up- and downstream of the coding region (Fig. 2A). This “knockout” plasmid was made using the MultiSite Gateway Three-Fragment Vector Construction Kit (Invitrogen) based on methods previously described (45). Two entry vectors were constructed to include 5’ and 3’ flanking sequences for recombination with the endogenous locus: pDONR_P2R-P3 contains the 1,875 bp sequence downstream of the AP2IX-4 genomic sequence and pDONR_P4P1R contains the 1,660 bp sequence upstream. A third entry vector, pDONR221, was constructed to contain the DHFR*-TS minigene. Each entry vector was constructed using the Gateway BP reaction, which consists of combining a PCR product with a pDONR vector; all primers used to generate these vectors are listed in Supplemental Table 1. The three entry vectors were combined in a Gateway LR reaction with destination vector pDEST_R4R3 to create the final knockout plasmids, which were then electroporated into RHQ or PruQ parasites. Pyrimethamine-resistant parasites were cloned by limiting dilution and examined by genomic PCR and RT-PCR to confirm the fidelity of each knockout (Fig. 2B).

### Complementation of the AP2IX-4 knockout

We complemented the PruQ*Δap2IX-4* parasites by restoring a cDNA-derived version of AP2IX-4 at the knockout locus and downstream of its endogenous promoter using single homologous recombination (Fig. 3A). A 2,116 bp sequence upstream of the start codon was amplified (primers shown in Supplemental Table I) and ligated to a cDNA-derived AP2IX-4 coding sequence using InFusion Cloning (Clontech) and integrated at the PacI restriction site into vector pLIC-3HA-HXGPRT. Vector pLIC-3HA-HXGPRT contains the HXGPRT drug selection cassette that replaced the DHFR*-TS minigene of its parental vector, pLIC-3HA-DHFR. Integration of the AP2IX-4 coding sequence into pLIC-3HA-HXGPRT fuses it to a C-terminal 3xHA tag. The resulting plasmid was electroporated into *Δap2IX-4* parasites, which were then selected in 25 µg/ml mycophenolic acid and 50 µg/ml xanthine and cloned by limiting dilution (46).

### RT-PCR

Total RNA was isolated from the designated parasites using TRIzol Reagent (Ambion) and treatment with DNaseI for 30 min at 37°C (Promega). The mRNA was reverse transcribed using Omniscript reverse transcriptase (Qiagen) and oligo(dT) primers as per manufacturer’s instructions; subsequent PCRs were performed using primers for AP2IX-4 or GAPDH (Supplemental Table 1).

### Western blot analysis

Immunoblotting of parasite lysate was used to detect AP2IX-4 in various clones by virtue of the 3xHA epitope tag. Parasite lysate was prepared in lysis buffer (150 mM NaCl, 50 mM Tris-Cl pH 7.5, 0.1% NP-40) and sonicated prior to resolving on SDS-PAGE and transfer to a nitrocellulose membrane. HA epitopes were detected using rat anti-HA antibody (Roche) at 1:5,000 dilution followed by secondary probing with horseradish peroxidase (HRP)-conjugated goat anti-rat antibody (GE Healthcare) at 1:2,000. To ensure equal loading of samples, we also probed with rabbit anti-β-tubulin antibody (kindly supplied by Dr. David Sibley, Washington University) at 1:2,000 dilution followed by HRP-conjugated donkey anti-rabbit antibody (GE Healthcare) at 1:2,000.

### Immunofluorescence assays (IFA)

HFF monolayers grown on coverslips in 12-well plates were inoculated with freshly lysed parasites. Infected monolayers were fixed in 3% para-formaldehyde (PFA) for 10 min, quenched in 0.1 M glycine for 5 min, and then permeabilized with 0.3% Triton X-100 for 20 min with blocking in 3% BSA/1xPBS for 30 min. *IFAs of tachyzoite cultures:* Rabbit polyclonal anti-HA primary antibody (Invitrogen) was applied at 1:2,000 dilution overnight at 4°C, followed by goat anti-rabbit Alexa Fluor 488 secondary antibody (Thermo Fisher) at 1:4,000 dilution. Detection of IMC3 was achieved using rat monoclonal anti-IMC3 antibody (kindly supplied by Dr. Marc-Jan Gubbels, Boston College) at 1:2,000 dilution overnight at 4°C, followed by goat anti-rat Alexa Fluor 594 antibody (Thermo Fisher) at 1:4,000 dilution. 4’,6-diamidino-2-phenylindole (DAPI, Vector Labs) was applied for 5 min as a co-stain to visualize nuclei. *IFAs of bradyzoite cultures:* Rat monoclonal anti-HA primary antibody (Roche) was applied at 1:1,000 dilution for 1 hour at room temperature, followed by goat anti-rat AlexaFluor 594 secondary antibody at 1:1,000 dilution. For the visualization of tissue cyst walls, FITC- or rhodamine-conjugated *Dolichos biflorus* lectin (Vector Lab) was applied at 1:1,000 dilution for 30 min at room temperature or 1:250 dilution overnight at 4°C, respectively. In IFAs co-stained for IMC3 or SAG1, rabbit polyclonal anti-HA primary antibody was applied at 1:2,000 dilution, followed by goat anti-rabbit Alexa Fluor 568 secondary antibody at 1:4,000 dilution. Detection of IMC3 was achieved using rat monoclonal anti-IMC3 antibody at 1:2,000 dilution overnight at 4°C, followed by either goat anti-rat Alexa Fluor 647 antibody or anti-rat Alexa Fluor 594 at 1:4,000 dilution. Detection of SAG1 was achieved using mouse monoclonal anti-SAG1 antibody (Abcam) followed by goat anti-mouse AlexaFluor 647 secondary antibody.

### RNA purification and microarray analyses

Total RNA was purified from intracellular PruQ, PruQ*Δap2IX-4*, or PruQ*Δap2IX-4*::AP2IX-4^HA^ parasites maintained at pH 7.0 (32-36 hours post-infection) or following 48 hours exposure to alkaline medium (pH 8.2) using the RNeasy Mini kit (Qiagen) according to manufacturer’s instructions. Two biological replicates were prepared for each parasite line under each experimental condition and RNA quality was assessed using the Agilent Bioanalyzer 2100 (Santa Cruz, CA). RNA samples were prepared for hybridization on the ToxoGeneChip as previously described (19). Hybridization data was analyzed using GeneSpring GX software package (v12.6.1, Agilent) and all datasets have been made available in the NCBI GEO database (GSE94045). Transcriptome data from previous publications was used to denote whether genes in our datasets were associated with expression in the bradyzoite stage (29, 47).

### Virulence and cyst burden in infected mice

To examine acute virulence, 5-6 week old female BALB/c mice were injected intraperitoneally with 10^5^, 10^6^, or 10^7^ parental PruQ, PruQ*Δap2IX-4*, or PruQ*Δap2IX-4*::AP2IX-4^HA^ parasites (8 mice per group). Plaque assays were performed for each sample and ensured equal viability between strains. Mice were examined daily and time to death was recorded. Serology performed on cardiac bleeds of infected mice confirmed presence of *Toxoplasma*. To assess cyst burden, mice were infected with 10^6^ parasites as described above and allowed to progress to chronic infection for 35 days (4 mice per group). Brains were then homogenized; homogenates were fixed, quenched, and permeabilized. Samples were blocked in 3% BSA/1xPBS/0.2% Triton X-100. To visualize cyst walls, rhodamine-conjugated *Dolichos biflorus* lectin (Vector Labs) was applied at 1:250 dilution overnight at 4°C. Cyst quantification was performed as previously described (48).

### Ethics statement

Mice were infected with *Toxoplasma* using an approved protocol (10852) from the Institutional Animal Care and Use Committee (IACUC) of the Indiana University School of Medicine (IUSM). The IUSM is accredited by the International Association for Assessment and Accreditation of Laboratory Animal Care.

## ACKNOWLEDGEMENTS

The authors thank Drs. David Sibley (Washington University) and Marc-Jan Gubbels (Boston College) for donating antibody reagents, Dr. Gustavo Arrizabalaga (Indiana University School of Medicine) for helpful discussions, and Gian Gballou for assisting with cyst burden quantification. We also thank Dr. Joanna Gress at the Functional Genomics Core Facility at Montana State University for help with hybridization of the ToxoGeneChip microarrays. This research was supported by Showalter Scholar Funding (to WJS) and the National Institutes of Health (AI089885 and AI124682 to MWW).

## SUPPLEMENTAL TABLES

**Supplemental Table 1. Primers used in this study.**

**Supplemental Table 2. Differentially expressed genes (FC≥2) under tachyzoite conditions in PruQ*∆ap2IX-4* compared to parental PruQ parasites.**

**Supplemental Table 3. Differentially expressed genes (FC≥2) under bradyzoite conditions in PruQ*∆ap2IX-4* compared to parental PruQ and *∆ap2IX-4*::AP2IX-4^HA^ parasites.**

